# PACIFIC: A lightweight deep-learning classifier of SARS-CoV-2 and co-infecting RNA viruses

**DOI:** 10.1101/2020.07.24.219097

**Authors:** Pablo Acera Mateos, Renzo F. Balboa, Simon Easteal, Eduardo Eyras, Hardip R. Patel

**Author notes:** Correspondence: Hardip R. Patel and Eduardo Eyras. Email addresses (PAM), (RFB), (SE), (EE), (HRP). These authors contributed equally to this work (Pablo Acera Mateos, Renzo F. Balboa).

## Abstract

Viral co-infections occur in COVID-19 patients, potentially impacting disease progression and severity. However, there is currently no dedicated method to identify viral co-infections in patient RNA-seq data. We developed PACIFIC, a deep-learning algorithm that accurately detects SARS-CoV-2 and other common RNA respiratory viruses from RNA-seq data. Using *in silico* data, PACIFIC recovers the presence and relative concentrations of viruses with >99% precision and recall. PACIFIC accurately detects SARS-CoV-2 and other viral infections in 63 independent *in vitro* cell culture and patient datasets. PACIFIC is an end-to-end tool that enables the systematic monitoring of viral infections in the current global pandemic.

## Background

Acute respiratory tract infections are the third largest global cause of death, infecting 545 million people and claiming 4 million lives every year (1–3). RNA viruses such as influenza, parainfluenza virus, respiratory syncytial virus, metapneumovirus, rhinovirus, and coronavirus are amongst the top pathogens causing respiratory infections and disease (4,5). Novel respiratory diseases, including coronaviruses, cross species boundaries repeatedly. Since December 2019, millions of people have been affected by COVID-19, an infectious zoonotic disease caused by severe acute respiratory syndrome coronavirus 2 (SARS-CoV-2, NCBI Taxonomy ID: 2697049). Novel zoonotic coronaviruses also caused the 2002-2003 outbreak of SARS-CoV respiratory disease with at least 8,098 known cases and the ongoing 2012-2020 outbreaks of Middle East respiratory syndrome coronavirus (MERS-CoV) with at least 2,519 known cases (5–8). This recurrent emergence of respiratory viruses warrants increased surveillance and highlights the need for rapid, accurate, and timely diagnostic tests.

Diagnostic testing, treatment, and disease severity are complicated by the occurrence of respiratory co-infections. Up to 40% of individuals infected with respiratory viruses test positive for co-infections with up to three different pathogens (9,10), and recent studies have reported that ∼20% of SARS-CoV-2 positive individuals had a co-infection with other respiratory viruses (11). Viral co-infections can alter the severity of disease and modify survival rates, and while COVID-19 remains poorly understood, early studies indicate a potential for increased mortality associated with influenza co-infections (12). Further studies are required to investigate the relationship between SARS-CoV-2 co-infections with prognosis and mortality rate (12,13).

Current diagnostic tests for respiratory infections are often limited in their capacity to detect co-infections. The current standard of viral detection for COVID-19 is based on polymerase chain reaction (PCR) assays directed towards SARS-CoV-2 (14), which do not detect co-infecting viruses. Multiple virus identification in clinical settings is typically performed by using multiplexed PCR assays with primers specifically designed to target known respiratory pathogens (15). However, this approach is generally used only with pathogens that are expected *a priori* and the range of pathogen detection is limited by the probe design. Additionally, these protocols must be updated as new species or strains are identified as clinically relevant.

High-throughput RNA sequencing (RNA-seq) provides an unbiased measurement of the RNA molecules present in a sample and can potentially enable the systematic detection of SARS-CoV-2 infections and co-infections. Multiple species identification has been effectively performed in sequence data in the context of metagenomics studies (16). Programs such as Kraken (17–19) use k-mers to taxonomically classify sequencing reads into species from metagenomic samples. However, these tools use large databases of species sequences to compare against, resulting in considerable storage and computing requirements. In contrast, machine learning based tools have the advantage of extracting required features and encapsulating the necessary information for sequence classification in a computationally efficient model. This approach has been successfully used in the past for sequence classification problems (20). For example, DeepMicrobes (21) uses deep learning for genus and species level classification of metagenomic DNA sequencing reads from human gut bacteria. Similarly, ViraMiner (22) uses a deep learning binary classifier to identify DNA viruses from human microbiome metagenomic reads. Despite these advances, there is currently no equivalent deep learning classifier for the detection of SARS-CoV-2 and possible co-infections by RNA viruses.

To address this limitation, we have developed PACIFIC, a deep learning model to detect the presence of SARS-CoV-2 and other common respiratory RNA viruses in RNA-seq data from patient samples. PACIFIC is an easy-to-use, streamlined tool useful for clinical and epidemiological applications in the context of the COVID-19 pandemic. Our tool accurately identifies and discriminates reads into five distinct classes: SARS-CoV-2, influenza (representing H1N1, H2N2, H3N2, H5N1, H7N9, H9N2, and Influenza B), metapneumovirus (representing 5 distinct assemblies), rhinoviruses (representing rhinovirus A and A1, B, C1, C2, C10 and other enteroviruses) and other coronaviruses (representing alpha, beta, gamma, and other unclassified coronaviruses). Extensive *in silico* tests show that PACIFIC achieves >99% precision, accuracy and recall. In addition, predictions in 63 infected human cell-lines and human primary samples demonstrate greater performance using PACIFIC for the detection of each virus class in comparison with alignment (BWA-MEM) and k-mer (Kraken2) based methods.

To the best of our knowledge, PACIFIC is the first software that uses deep learning to classify different RNA viruses from RNA-seq reads. By enabling the systematic identification of co-infections, we anticipate that PACIFIC can aid the clinical management of COVID-19 patients during the current pandemic and the surveillance of respiratory infections in future epidemiological studies.

## Results

### PACIFIC model

PACIFIC is a deep learning method designed to classify RNA-seq reads into five distinct respiratory virus classes and a human class (Figure 1a). The model architecture for PACIFIC is composed of an embedding layer, a convolutional neural network, and a bi-directional long short-term memory (BiLSTM) network that ends in a fully connected layer (Figure 1b). One of the main advantages of deep neural networks compared to other machine learning models in the context of sequence classification is the ability to extract relevant complex classification features from DNA or RNA sequences without having to explicitly define them *a priori*. However, the strategy to encode nucleotide sequences must be carefully considered, as it can dramatically affect the performance of the classifier (23). PACIFIC implements an embedding layer, which boosts the performance of the model in comparison with other encoding approaches (24). PACIFIC first converts nucleotide sequences into k-mers, assigns them to numerical tokens and converts these tokens into dense representations using a continuous vector space.

**Figure 1.**
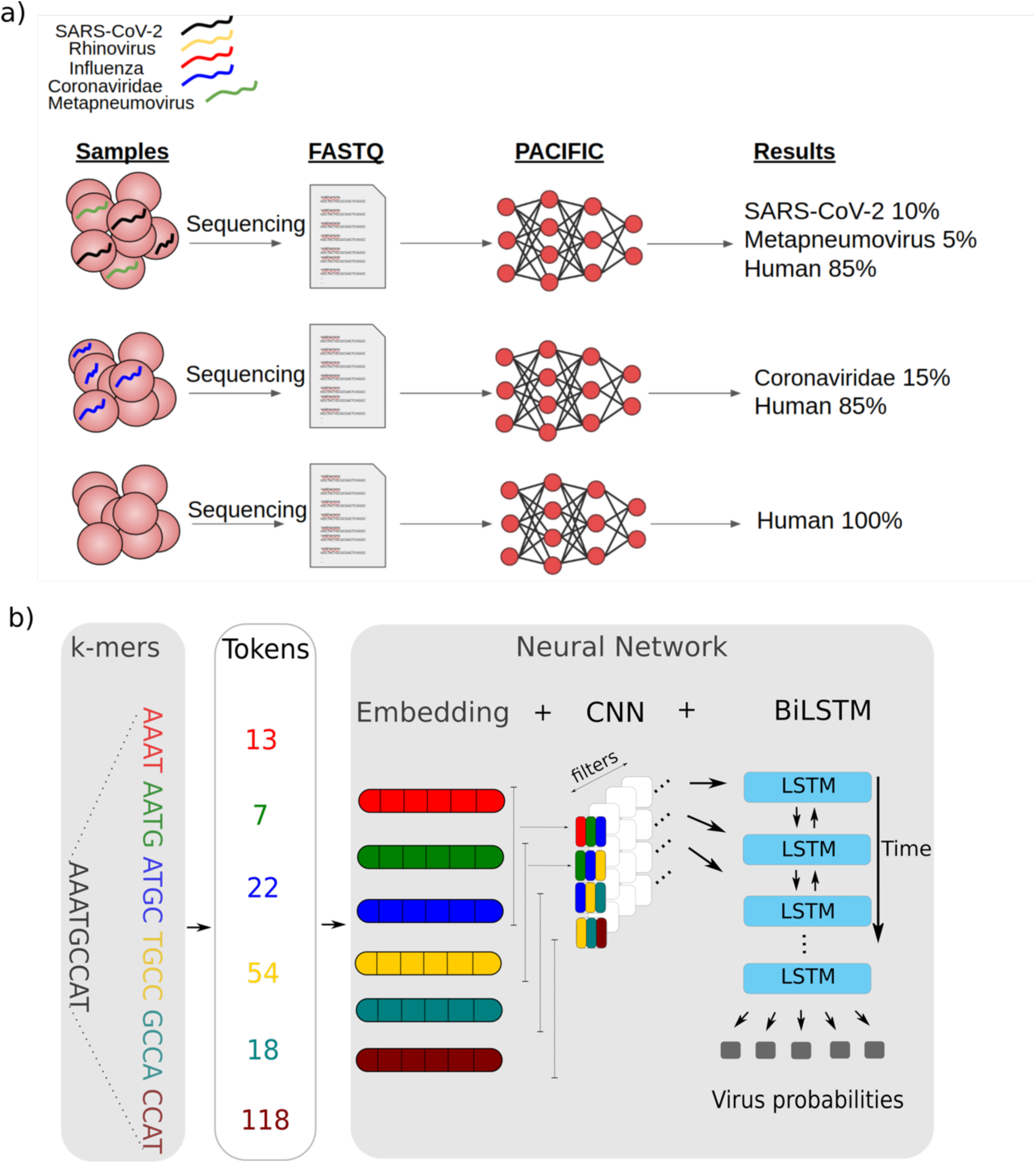
Overview of PACIFIC and its model architecture. **a)** PACIFIC uses FASTQ or FASTA files as inputs to make read-level predictions and report the relative percentage of RNA virus and human reads in a sample. **b)** Schematic view of PACIFIC deep neuronal network architecture. PACIFIC uses embedding, convolutional neural network and BiLSTM layers. The model is trained using *in silico* generated sequences from RNA virus genomes and the human transcriptome.

The use of a convolutional neural network adds several advantageous properties to the model. One advantage is location invariance (25), which allows the model to identify combinations of features with predictive value regardless of their relative position along the sequence. In addition, each filter used in the convolution layers can capture the predictive value of specific regions or combinations of k-mers.

After the convolution layers, PACIFIC uses a pooling layer to decrease the dimensionality of the feature space while maintaining essential information. PACIFIC uses a BiLSTM to model long-range dependencies in nucleotides, which provides the capacity to incorporate complex relationships in the input sequence that are sometimes ignored by single LSTMs (26). PACIFIC then implements a dense layer to estimate posterior probabilities for each of the five classes considered: Coronaviridae, Influenza, Metapneumovirus, Rhinovirus, and SARS-CoV-2. We included a sixth class in the model (Human) to classify RNA-seq reads derived from the human host. Finally, PACIFIC takes the reverse-complement of each predicted read, and only assigns the read to a particular class if the posterior probabilities of both the forward and reverse-complemented versions of the read for that class are ≥ 0.95.

### Properties of PACIFIC training data

PACIFIC was trained using 7.9 million 150nt long random fragments from 362 viral genome assemblies belonging to one of five viral classes (SARS-CoV-2, Influenza, Metapneumovirus, Rhinovirus and Coronaviridae) and the human transcriptome (Additional file 1: Table S1) (Methods). *In silico* fragments from both strands were generated without errors to accommodate paired-end sequenced reads and to retain the natural variation between genomes in each class. We used 90% of the data for training and 10% to tune the hyperparameters and network architecture.

The selection and grouping of virus classes were based on several considerations. First, we wanted PACIFIC to accurately detect SARS-CoV-2 as an independent class and to discriminate it from other coronaviruses. Second, we selected viruses that have been recently reported to appear as co-infections with SARS-CoV-2 (11). Third, we restricted our selection to viruses for which humans have been defined as one of the host species in NCBI Taxonomy database (27). Fourth, as the majority of reads in a sample are expected to be derived from human RNAs, we included an independent human class representing the human transcriptome to avoid the misclassification of human reads as viral origin.

### K-mer length selection and sequence divergence

As input reads are divided into k-mers within the model, we investigated appropriate virus and human k-mer properties. A k-mer length of 9 was previously reported to be the optimal k-mer length for the phylogenetic separation of viral genomes (28). However, 9-mer profiles of SARS-CoV-2 and the human transcriptome have not been previously explored. We computed all-vs-all Jensen-Shannon divergence (JSD) scores using 9-mers to confirm that k=9 is the effective *k*-length to distinguish between the six PACIFIC classes. JSD is a symmetric measure of (dis)similarity that accounts for shared k-mer frequency distributions between a pair of sequences (29). JSD values range between 0 for identical sequences and 1 for two sequences that do not share any k-mer. Overall, inter-class JSD values were higher compared to the intra-class JSD values for 9-mers, which confirm that 9-mers are effective at separating sequences belonging to different viral classes and human transcripts (Additional file 2: Figure S1). Specifically, the average JSD between the SARS-CoV-2 class and the Coronaviridae class was 0.786, which was greater than 0.767 intra-class JSD for the Coronaviridae and 0.002 for the SARS-CoV-2 class, thus indicating sufficient divergence for their separation into distinct classes. Given these results, we decided to encode input sequences as 9-mers with a stride of 1.

### PACIFIC testing shows high precision and recall for simulated data

Performance metrics (false positive rates (FPRs), false negative rates (FNRs), precision, recall, and accuracy) were calculated using *in silico* generated reads that modelled sequencing-induced substitution and indel errors from sample mixtures with known class labels. We generated 100 independent datasets of 150nt single end reads with Illumina HiSeq2500 errors using ART (30). Each dataset contained ∼700,000 reads and was comprised of approximately 100,000 reads from each of the 6 classes in the PACIFIC model, plus ∼100,000 reads from unrelated viral genomes (Methods).

First, we compared the performance metrics between predictions using only the forward strand of a read (*single prediction*) and predictions using both the forward and reverse-complemented strands (*double prediction*) (Figure 2). In *single prediction*, a read was assigned the class label with the highest posterior probability for the forward sequence if the posterior probability for a class was ≥ 0.95. For *double prediction*, the read was predicted to be of a given class if the posterior probability of the predicted forward and reverse-complemented read was ≥0.95 and predictions agreed on the same class. The average FNR across 100 datasets in *double prediction* relative to *single prediction* increased by 1.45× for Coronaviridae, 1.49× for Influenza, 1.41× for Metapneumovirus, 1.40× for Rhinovirus and 1.62× for SARS-CoV-2 (Figure 2). However, we observed a large decrease in the FPR in *double prediction*. The average FPR in the 100 datasets decreased by 4.60× for Coronaviridae, 11Confusion matrix using SARS-CoV-2 as.02× for Influenza, 16.55× for Metapneumovirus, 3.92× for Rhinovirus and 16.47× for SARS-CoV-2 class in *double prediction* relative to *single prediction* (Figure 2). Concomitantly, the average precision for all viral classes increased and the average recall decreased by a small margin. Due to these observations, *double prediction* was implemented as the standard classification approach in PACIFIC.

**Figure 2.**
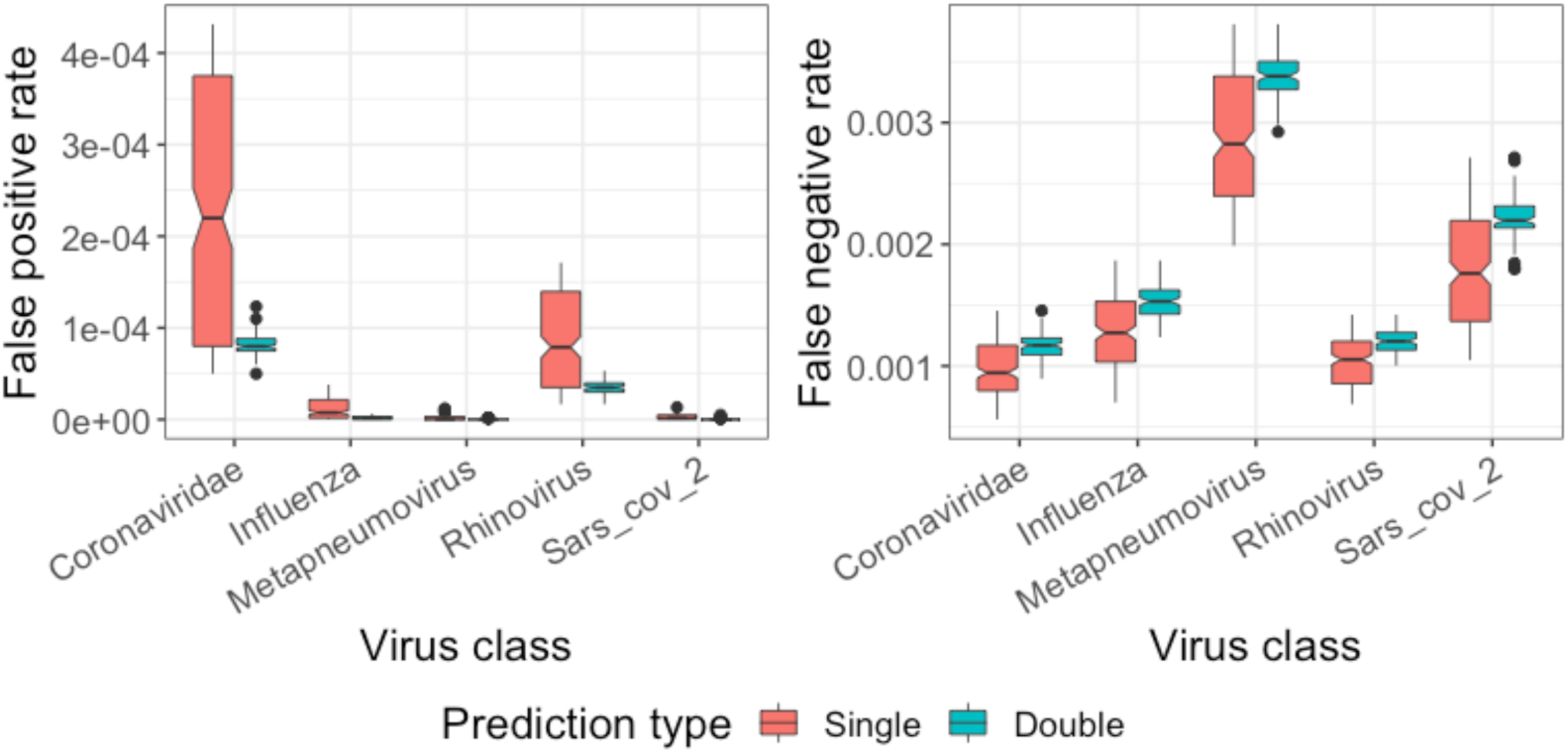
Comparison of false positive and false negative rates between *single* and *double predictions*. *Single prediction* (red) results in relatively higher false positives and lower false negatives compared to *double prediction* (green) where predictions are made on the forward strand of a sequence and its reverse complement.

Overall, PACIFIC achieved high precision (average ≥ 0.9995), recall (average ≥ 0.9966) and accuracy (average ≥ 0.9995) for each of the virus classes (Table 1). For the human class, the average precision was lower at 0.50140 compared to the viral classes; this is attributed to the large number of reads (>99%) from unrelated viral genomes being assigned to the human class. As a result, sequences that do not belong to any of the viruses in the model are unlikely to be mislabelled as one of the virus classes.

**Table 1.**
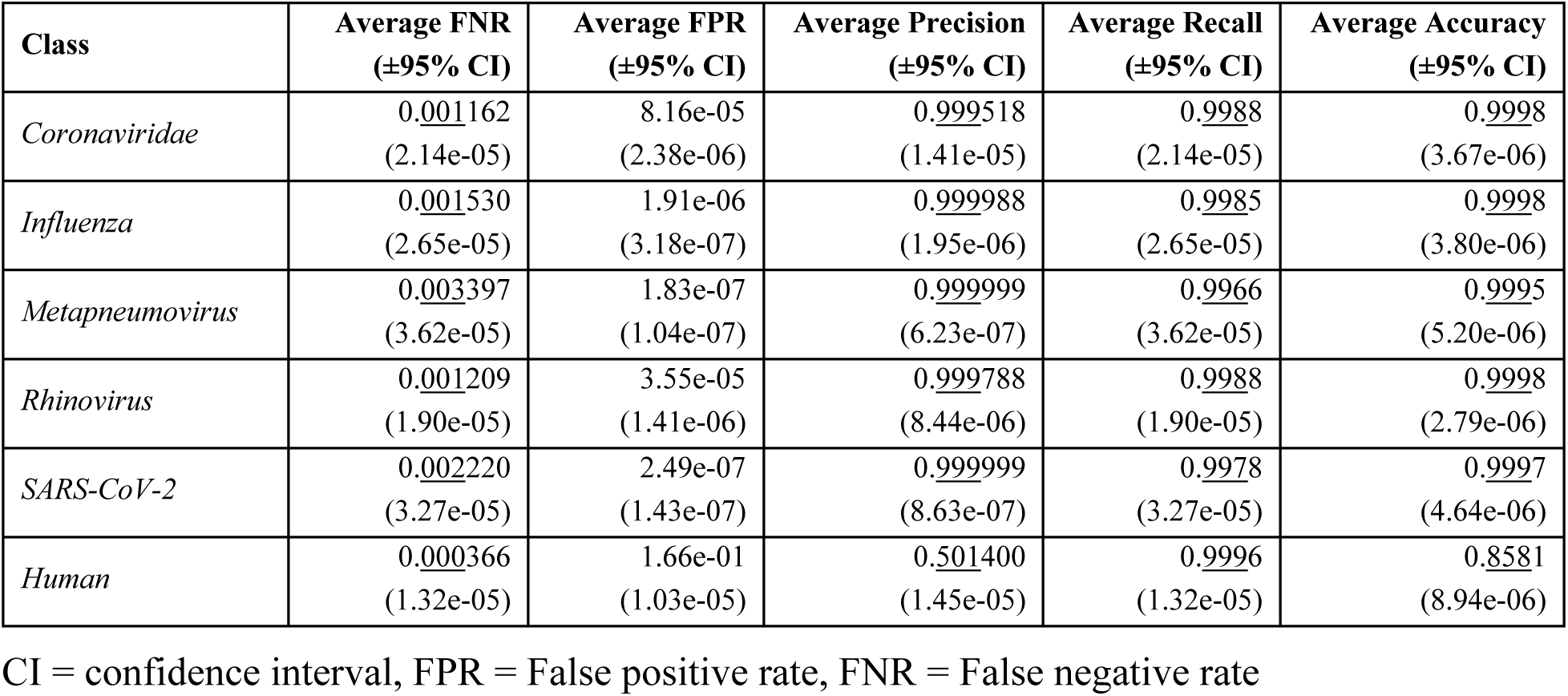
PACIFIC performance metrics for each class in 100 independent simulated datasets.

Using the same *in silico* datasets, we then assessed the effect of mismatches on FPR and FNR. All 100 controlled datasets contained ∼22% of reads with substitutions and indel errors relative to the reference genomes. FNRs were higher for mismatch-containing reads relative to exact reads for all viral classes, increasing 5 to 64-fold (Figure 3). In contrast, FPRs increased 0.98 to 2-fold for mismatch-containing reads for all classes. These results suggest that FPRs are relatively less affected by the presence of mismatches compared to FNRs for viral classes. Despite relative differences of FPR and FNR in mismatch-containing reads compared to exact reads, PACIFIC achieved high precision (0.9994), high recall (0.9909) and high accuracy (0.9982) for mismatch-containing reads for all five viral classes (Additional file 1: Table S2).

**Figure 3.**
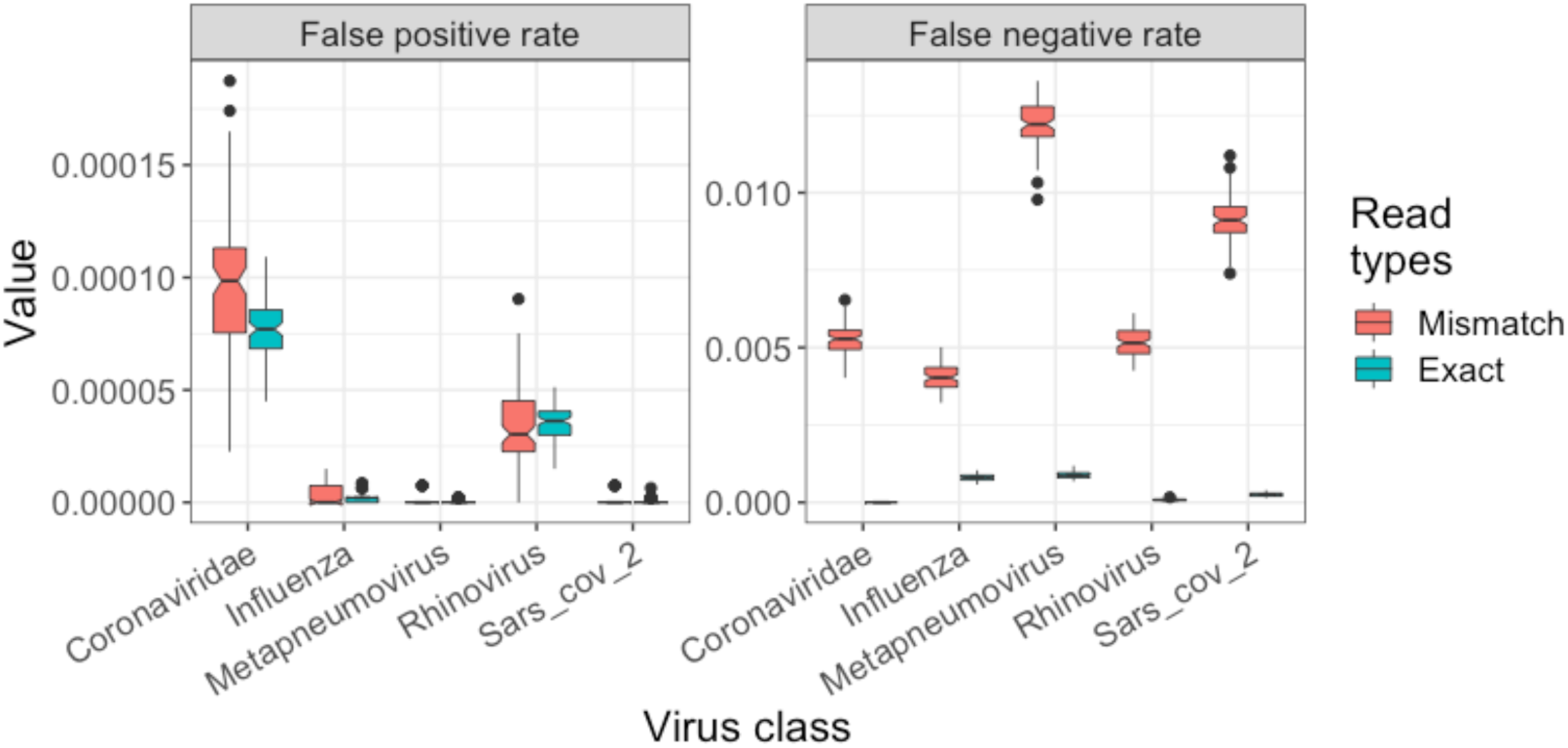
False positive (left panel) and false negative (right panel) rates for reads identical to the corresponding reference genome (Exact, green), and for reads with mismatches with respect to their reference genome (Mismatch, red).

Reads derived from unrelated virus genomes contributed most of the false positives for all predicted classes (Figure 4). Of note, there was negligible cross contamination between viral class labels. The largest inter-viral class misclassification was from SARS-CoV-2 to Coronaviridae, where out of 100,000 SARS-CoV-2 reads, ∼2.9 were misclassified as Coronaviridae. Other inter-viral class misclassifications were between 0-0.2 reads. The majority of false negatives for each viral class were either discarded because of the *double prediction* criteria (rc_discarded in Figure 4), or because they did not meet the minimum 0.95 posterior probability criteria (pDiscarded in Figure 4). Taken together, our results demonstrate that PACIFIC is highly specific and sensitive for all five viral classes, with negligible false positive and false negative rates.

**Figure 4.**
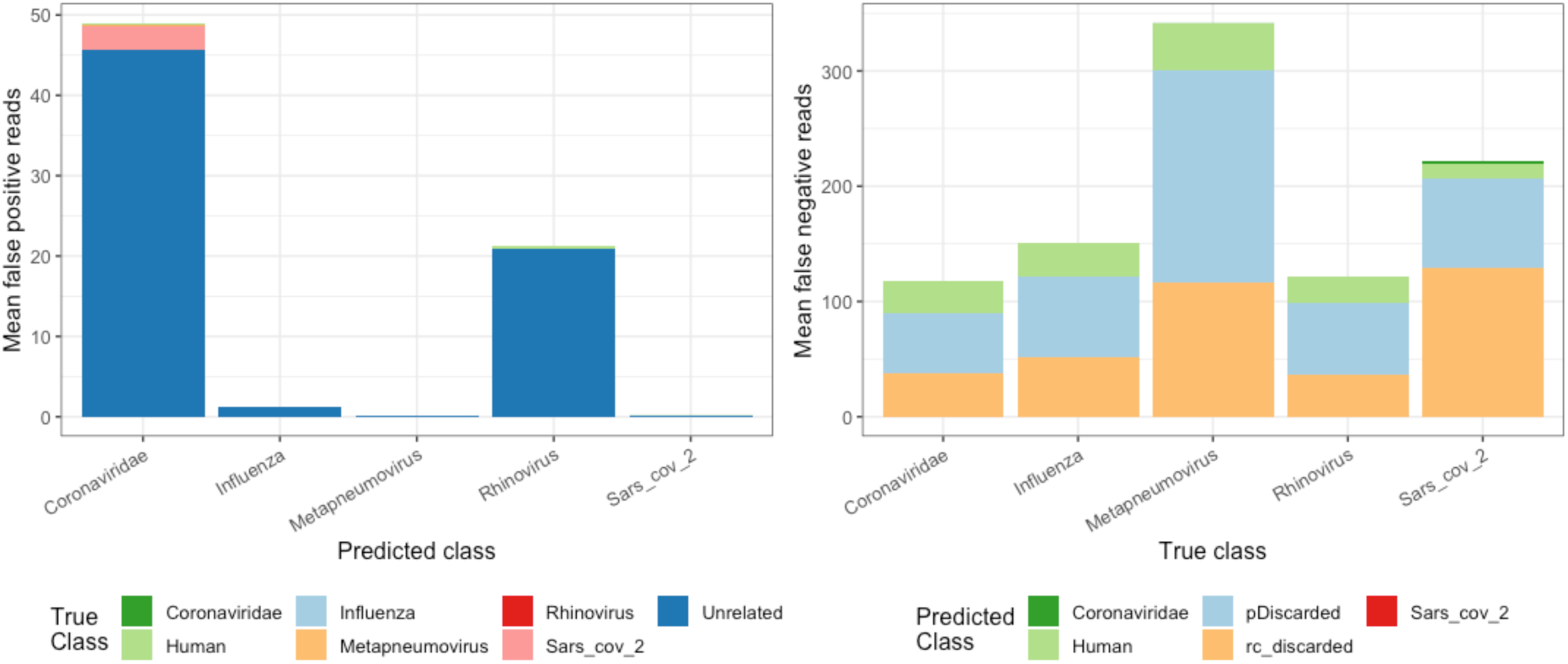
False positive and false negative assignments for 100 independently simulated datasets. Left panel - Average number of false positives (y axis). True labels are indicated by colour for each class and predicted labels are given in the x-axis. Right panel - Average number of false negatives (y axis). Predicted labels are indicated by colour for each class and true labels as indicated in the x-axis. pDiscarded - reads that do not reach the 0.95 posterior probability cut-off; rc_discarded - reads that were discordantly predicted using *double prediction*.

### Establishing virus detection thresholds

In the previous section, we showed that PACIFIC displayed low FPR in balanced datasets with similar proportions of reads from each class. However, incorrect predictions about the presence or absence of a virus in a sample could lead to misguided follow-ups and the unnecessary use of valuable clinical resources. Therefore, we decided to establish the minimum percentage of reads for each viral class required to confidently predict the presence of a virus in a sample.

In practice, RNA-seq data will be unbalanced with almost all reads originating from the human transcriptome mixed with variable proportions of viral reads. To model this imbalance in class proportions, we simulated 500 independent datasets (100 for each class), each containing 500,000 150nt long reads using ART (30) with Illumina HiSeq2500 error profiles. Each dataset contained variable proportions of simulated reads for 4 of the 5 viral classes, plus human and unrelated viral genomes. One of the five viral classes was intentionally excluded, and the excluded class was considered to be the test class. All reads assigned to this test class were counted as false positives. PACIFIC achieved similar average FPRs to the benchmarking experiments using balanced datasets (Table 2).

**Table 2.** Average false positive reads across 100 experiments for each viral class.

| Class | Average FP%<br>(Balanced) | Average FP%<br>(Unbalanced) | FP % threshold* |
| --- | --- | --- | --- |
| Coronaviridae | 0.00816 | 0.006664 | 0.0405 |
| Influenza | 0.000191 | 0.000062 | 0.000807 |
| Metapneumovirus | 0.0000183 | 0.000006 | 0.000154 |
| Rhinovirus | 0.00355 | 0.005392 | 0.0418 |
| SARS-CoV-2 | 0.0000249 | 0.000010 | 0.000213 |

FP = False positive, Balanced = 100 experiments with equal proportion of reads from each class, Unbalanced = Variable proportion of reads from all classes and no reads from the test class. \* These thresholds represent 0.99 quantile for FP% for that class.

Assessment of the distribution of false positive rates for each viral class and skewness-kurtosis plots indicated that the percentage of false positives observed in unbalanced datasets followed a Beta distribution. Therefore, we used moment matching to estimate the shape parameters for the quantile function of the Beta distribution and determined the numeric threshold above which 99% of false positive samples were excluded. Using these thresholds, a sample would be classified as positive for Coronaviridae if >0.0405% reads were labelled as Coronaviridae by PACIFIC. Similarly, these limits were >0.000807% for Influenza, >0.000154% for Metapneumovirus, >0.0418% for the Rhinovirus, and >0.000213% for the SARS-CoV-2 class (Table 2).

### PACIFIC accurately detects viruses in human RNA-seq samples

Next, we assessed PACIFIC’s performance in classifying viral reads in RNA-seq data derived from human biological samples and compared its output with alignment-based (BWA-MEM) and k-mer based (Kraken2) approaches. To reduce bias in the comparisons, we built the BWA-MEM index and Kraken2 database using the same virus genome assemblies and the human transcriptome that were used for PACIFIC training (Additional file 1: Table S1). Additionally, we used the same percentage thresholds determined in the previous section (Table 2) for all three methods to assign the presence of a virus in a sample. All three methods were applied to 63 human RNA-seq datasets from independent research studies, with five of them known to contain SARS-CoV-2 (31,32). Four RNA-seq datasets were derived from primary human lung epithelium cells (NHBE) infected with SARS-CoV-2 *in vitro* (NCBI SRA accession: SRX7990869) (31) and one dataset was from a patient bronchoalveolar lavage fluid sample (NCBI SRA accession: SRR10971381) that was positive for SARS-CoV-2 (32). In addition, we analysed RNA-seq datasets for 48 airway epithelial cell samples from the GALA II cohort study, of which 22 were reported to contain respiratory infection viruses (33,34), and 10 were reportedly devoid of infections by the viral classes studied here, as indicated by the sample metadata and the corresponding publications (Additional file 1: Table S3).

For the five samples that were positive for SARS-CoV-2, PACIFIC assigned 0.047%-0.048% of all reads to the SARS-CoV-2 class for the *in vitro* infected cells and 0.19% of all reads in the patient sample; all five samples were above the established detection threshold for that class (>0.000213%) (Figure 5). Similarly, BWA-MEM and Kraken2 successfully identified SARS-CoV-2 reads above the detection threshold in these five samples (Figure 5a). All three methods accurately predicted the presence of SARS-CoV-2 and the absence of other virus classes in these samples as per class specific detection thresholds.

**Figure 5.**
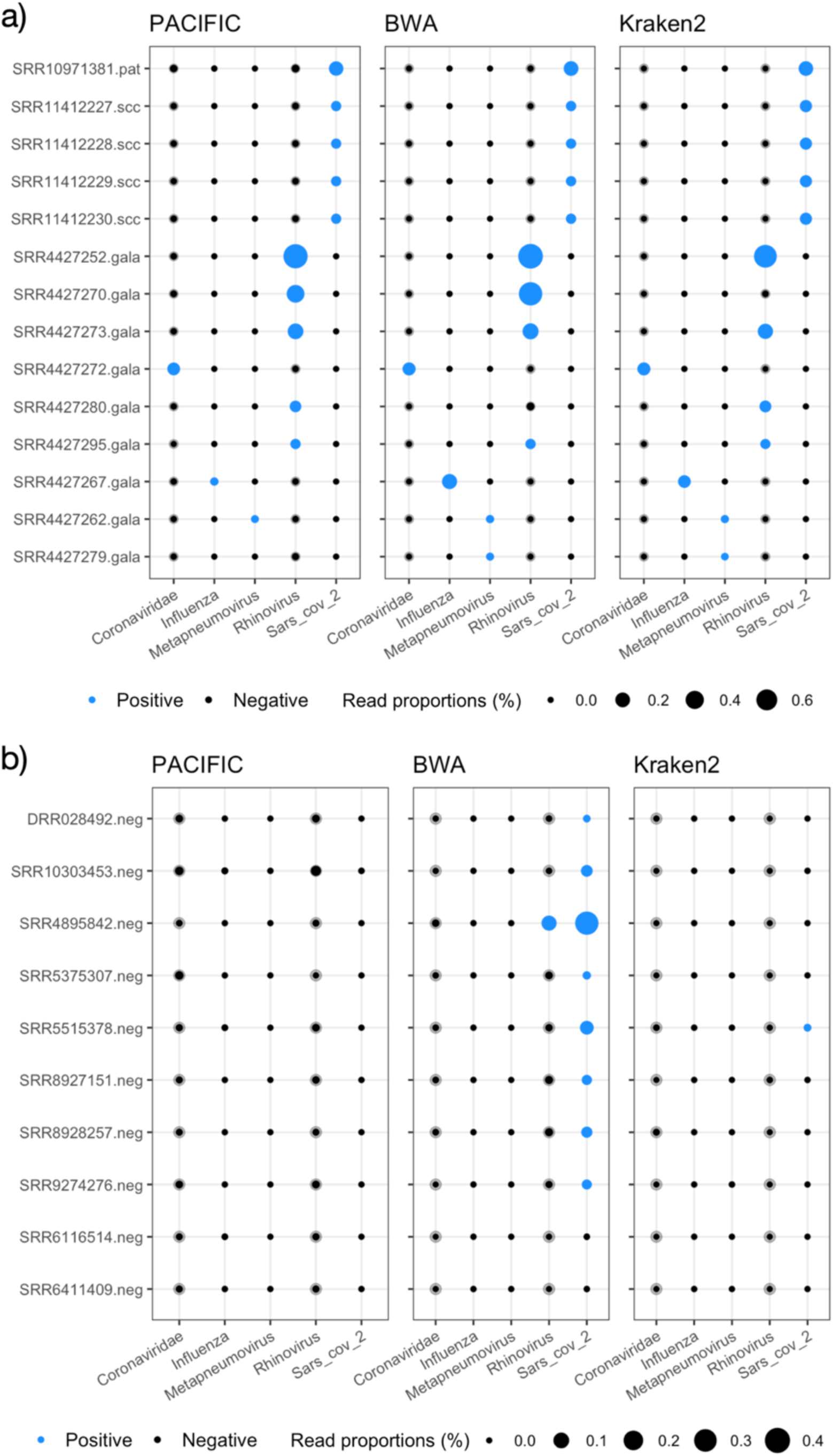
PACIFIC, BWA-MEM and Kraken2 virus predictions in RNA-seq data. Transparent grey filled circles represent detection thresholds for each class, overlaid with black filled circles representing the percentage of predicted reads using PACIFIC (left panel), BWA-MEM (centre panel) and Kraken2 (right panel). Circles are filled blue when the percentage of reads for a class are above detection thresholds described in Table 2. **(a)** RNA-seq samples predicted to be positive for at least one viral class by PACIFIC, BWA-MEM, or Kraken2. Samples include a SARS-CoV-2-infected human patient bronchoalveolar lavage fluid sample (.pat), four *in vitro* SARS-CoV-2-infected NHBE cell lines (.scc) and 9 samples from the GALA II cohort (.gala). **(b)** Human RNA-seq samples without expected viral infections (.neg; n=10). Samples were selected from the NCBI SRA database without any evidence of any infection.

We subsequently tested the 48 samples from the GALA II cohort (33,34). Of these, 22 were reported to contain between 4 and 164,870 reads from respiratory viruses (human rhinovirus, respiratory syncytial virus, human metapneumovirus or human parainfluenza viruses I, II and III). PACIFIC, BWA-MEM and Kraken2 identified 9 samples as positive for one of the five virus classes considered, and 39 samples as negative for the same viral classes (Figure 5a; Additional file 2: Figure S2, Additional file 1: Table S3). The discrepancy between these results and the original study (34) could be partly explained by the exclusion of the Respiratory Syncytial Virus class in our analyses. However, further verifications could not be performed because sample labels provided in the manuscript were mismatched with the submitted sequence data (see correction (35)).

From the 9 positive samples, six were concordantly labelled positive for the same virus class by all three methods: three samples were positive for Rhinovirus, and one for Coronaviridae, Influenza, and Metapneumovirus, respectively (Figure 5a). In contrast, the other three samples were discordantly labelled by one of the three methods. One sample (SRR4427279) was classified as positive for Metapneumovirus by BWA-MEM and Kraken2 but not by PACIFIC. BWA-MEM and Kraken2 collectively assigned 28 reads to the Metapneumovirus class as opposed to 0 reads by PACIFIC. To investigate the origin of these reads, we used BLASTN searches against the NCBI nucleotide *(nt)* database encompassing sequences from all domains of life and extracted the best hit for each read (Methods). All 28 reads had their best hits to human respiratory syncytial virus A sequences (E-values ≤ 4.07e-68; bit-scores ≥ 267). Further analysis showed that metapneumovirus was not identified in any of the top 10 significant hits (Additional file 2: *Extension of BLAST analysis*). Another discordant sample (SRR4427270) was positive for the Rhinovirus class by PACIFIC (1,065 reads) and BWA-MEM (2,338 reads) but not by Kraken2 (3 reads). BLASTN searches showed that best hits (3,145/3,150 total BLAST alignments) were sequences from one of Enterovirus C105, Enterovirus C or Human enterovirus C105 (E-values ≤ 9.46e-41, bit-scores ≥178). Rhinoviruses and other enteroviruses are taxonomically part of the Enterovirus genus (36,37). The final discordant sample (SRR4427280) was labelled positive for the Rhinovirus class by PACIFIC (355 reads) and Kraken2 (387 reads) but not by BWA-MEM (29 reads) using our thresholds (Figure 5a). BLASTN searches revealed that the majority of reads (385/390 collectively between PACIFIC and Kraken2) had their best hits to Rhinovirus C (E-values ≤ 4.47e-53; bit-scores ≥ 219). BWA-MEM therefore failed to assign most Rhinovirus reads classified by the other two methods.

In addition to the 53 samples tested above, we analysed 10 publicly available human RNA-seq datasets without expected viral infections using all three methods. In particular, all ten samples were registered in the NCBI SRA database on or before 17 October 2019 and therefore were unlikely to contain SARS-CoV-2 (Additional file 1: Table S3). Of note, two samples (SRR8927151 and SRR8928257) were published in February 2020 (Additional file 1: Table S3) (38). However, the two NCBI BioProjects for these samples were registered on 18 April 2019.

PACIFIC accurately predicted all 10 samples as negative for viral infections using our detection thresholds. In contrast, Kraken2 assigned one sample (SRR5515378) as positive for SARS-CoV-2 with 256 reads assigned (Figure 5b). Further verification with BLASTN searches confirmed that 242 out of the 256 reads mislabelled by Kraken2 aligned to the Mycoplasma bacterial genus (E-values ≤2.76e-23, bit-scores ≥121), indicating false positive assignments by Kraken2 for these reads.

BWA-MEM performed relatively worse in these 10 datasets, with 8 samples classified as positive for SARS-CoV-2 (Figure 5b). A total of 15,395 reads were aligned to SARS-CoV-2 genomes from these 8 samples, ranging from 86 to 6,901 reads in any given sample. BLASTN searches of these SARS-CoV-2 assigned reads showed *Homo sapiens* as the best hit (E-values ≤ 5.78e-07, bit-scores ≥ 65.8) for 4,352 (28%) reads. Further examination revealed that 53% of all 9-mers in these reads were poly-A or poly-T derived, suggesting low-complexity sequences. In addition, SRR4895842 was also assigned as co-positive for the Rhinovirus class in addition to SARS-CoV-2 by BWA-MEM (Figure 5b). BLASTN searches of 406 reads assigned to Rhinovirus within this sample revealed that 303 reads had best hits to *Homo sapiens*, and 48 reads had their best hits to *Pan paniscus* (E-value ≤ 6.06e-10; bit-scores ≥ 76.8). These reads had 19% of their 9-mers derived from poly-A or poly-T sequences, suggesting low-complexity sequences.

Overall, these results show that PACIFIC can accurately identify viral reads and the use of our detection thresholds assisted in correctly establishing the presence or absence of viral classes in RNA-seq data from biological samples with better accuracy than existing methods.

## Discussion

We have developed PACIFIC, a deep learning-based tool for the detection of SARS-CoV-2 and other common respiratory viruses from RNA-seq data. To the best of our knowledge, PACIFIC is the first deep learning model that performs detection of SARS-CoV-2 and different RNA virus groups using short-read sequence data with >0.99 precision, recall and accuracy. A recent analysis of 4,909 scientific articles identified 47 models for detecting COVID-19, 34 of which were based on medical images (39). This study concluded that these predictive models were, in general, poorly described and contained multiple biases, likely resulting in unreliable predictions when applied in practice. To overcome these potential limitations, we used multiple diverse and independent simulated datasets reflecting realistic scenarios to validate the performance of PACIFIC. Importantly, PACIFIC was successfully applied to 63 RNA-seq datasets derived from infected cell cultures and patient samples for the detection of viral infections, demonstrating that PACIFIC can be applied to human-derived RNA-seq datasets and assist in clinical settings.

In 2013, the World Health Organisation launched the Battle against Respiratory Viruses (BRaVe) initiative, which identified six research strategies to tackle and mitigate risks of death due to respiratory tract infections. One of the proposed strategies was to “*improve severe acute respiratory infection diagnosis and diagnostic tests amongst others”* (40). High-throughput sequencing-based approaches can provide immense diagnostic potential and facilitate molecular epidemiological studies, thereby contributing towards the BRaVe initiative’s goals (41,42). It is more important than ever to explore and determine the diagnostic potential of RNA-seq for the SARS-CoV-2 pandemic.

A comprehensive study using multiplex RT-PCR and a sequencing-based metagenomic approach revealed that RNA-seq has sufficient sensitivity and specificity to be applicable in the clinic for respiratory viruses (42). However, the use of RNA-seq in diagnostic settings is often complicated due to complex analytical workflows (34,42). A typical workflow for virus detection in high-throughput sequencing data involves quality assessment and filtering of raw data, removal of host sequences, *de novo* assembly of remaining reads, and lastly, the alignment and annotation of the generated contigs (43). Implementation of these workflows require expert knowledge of bioinformatics software and databases and often dedicated computing facilities. PACIFIC overcomes these limitations by modelling the differences in k-mer content of respiratory viruses and human sequences in a model that is efficient in compute and storage requirements, easy to use, and therefore applicable in contexts with minimal resources. Specifically, we have designed PACIFIC to be run as a single command using raw RNA-seq data as the only required input to obtain quantified predictions about viral classes within a sample.

Despite the higher costs of sequencing compared to PCR-based experiments, multiplexing, block-testing or pooling strategies (44) could be implemented for unbiased cost-effective testing. For example, sequencing with Illumina platforms could be done with 96 samples per lane using multiplexing, reducing the sequencing cost per sample. In this scenario, the number of reads obtained per sample could be approximately 200,000 or higher. We have demonstrated the accuracy of PACIFIC in a variety of sample sizes, which suggests the potential value of this approach.

One of the major challenges in the identification of virus classes is the high rate of natural sequence variation for RNA viruses (45,46), in addition to high-throughput sequencing induced errors and artefacts, and the presence of low-complexity A-rich sequences common to the host transcriptome. We showed that 22% of reads containing mismatches and indel errors were accurately assigned to a virus class by PACIFIC with negligible loss in sensitivity at a sample level. Given the ability of PACIFIC to accurately assign error-containing reads, we speculate that PACIFIC is applicable to cases where viruses present natural sequence variation. In such cases, or when new species are required to be added to the model, strategies like transfer learning can be used to update the model without the need to retrain the entire model, with low computational cost (47). Future versions of PACIFIC could focus on training a model that incorporates class specific mutation rates and sequence diversity to reduce the need for regular updates as new viral mutations emerge.

PACIFIC is intentionally focused on the identification of viral classes reported to be co-infecting along with SARS-CoV-2 (12). Therefore, samples containing other viruses and bacterial infections may require additional analysis. Future versions of PACIFIC could include the classification of a broader range of virus and bacterial classes at a species level, and variable input read lengths to increase PACIFIC’s utility in other contexts.

## Conclusions

PACIFIC is a powerful end-to-end and easy to use tool that predicts the presence of SARS-CoV-2, Influenza, Metapneumovirus, Rhinovirus and other Coronavirus class-derived sequences directly from RNA-seq data with high sensitivity and specificity. PACIFIC will enable effective monitoring and tracking of viral infections and co-infections in the population in the context of the COVID-19 global pandemic and allow for the development of new strategies in molecular epidemiology of co-infections to understand variable host responses and improve the management of infectious diseases caused by viruses.

## Methods

PACIFIC and other associated software written for this manuscript is available at https://github.com/pacific-2020/pacific. We have used Python (version 3), scipy (v1.4.1), numpy (v1.18.1), scikit (v0.23.1), pandas (v1.0.1), tensorflow (v2.2.0), keras (v2.3.1), R (v3.6), tidyverse (v1.3.0), Biobase (v2.46.0) and Perl (v5.26) in our analysis.

### Training data

We downloaded 362 virus genomes from the NCBI assembly database corresponding to five classes of single stranded RNA viruses (Table 4, Additional file 1: Table S1). GenBank assembly identifiers and assembly versions with other metadata are listed in Additional file 1: Table S1. Since our focus was to detect co-infections with SARS-CoV-2, we made a separate class for SARS-CoV-2 containing 87 different assemblies (Table 4). The *Coronaviridae* class contained 12 genomes of alpha, beta, gamma and unclassified coronaviruses. The Influenza class contained assemblies of influenza A (H1N1, H2N2, H3N2, H5N1, H7N9, and H9N2 strains) and influenza B viruses. For the Rhinovirus class, assemblies of rhinovirus A (including A1 strain), B, C (including C1, C2, and C10 strains), and unlabelled enterovirus were grouped together. There were five distinct assemblies for metapneumovirus which were grouped into a single class. We included Human GENCODE (48) canonical transcript sequences (downloaded from Ensembl v99 database (49)) as an additional class to distinguish sequencing reads derived from the human transcriptome. We generated between 0.44 and 3.5 million 150nt-long fragments *in silico* for each class using a custom Perl script available at https://github.com/pacific-2020/pacific (generatetestdata.pl, Table 4). These training sequences were randomly sampled without any base substitutions and were derived from both strands of the genome assemblies.

**Table 4.**
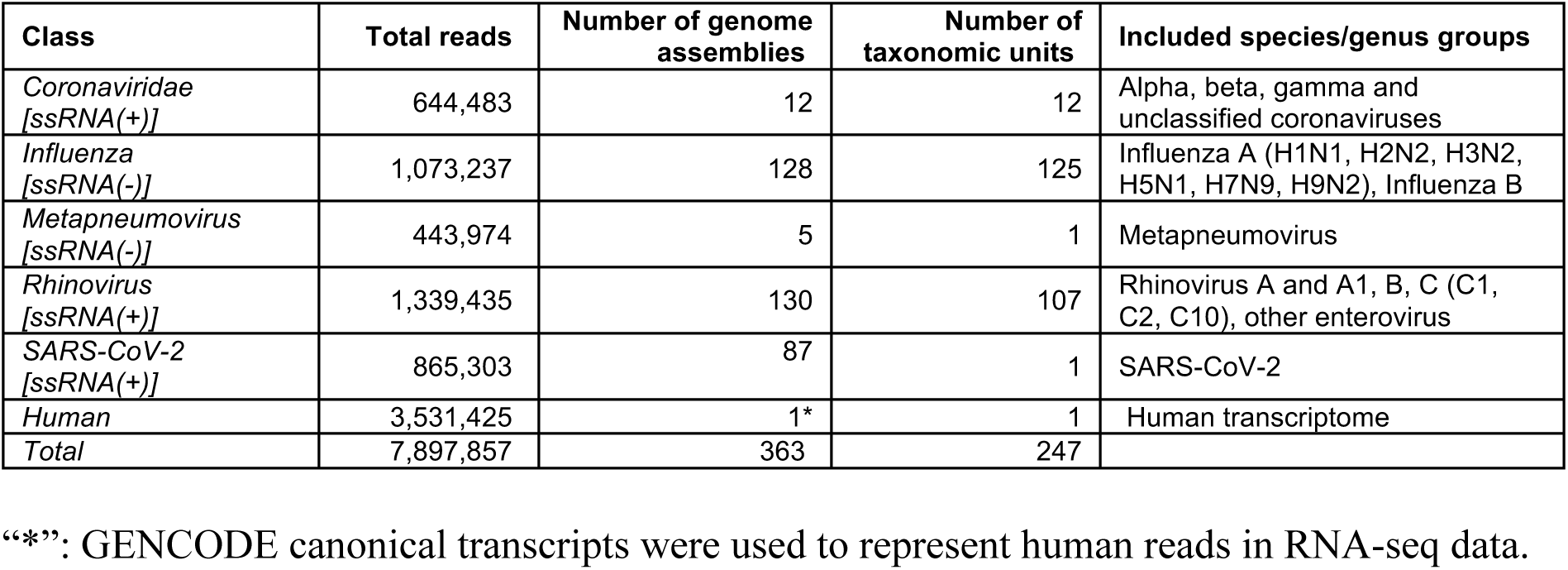
Summary of training classes used for PACIFIC.

| Class | Total reads | Number of genome assemblies | Number of taxonomic units | Included species/genus groups |
| --- | --- | --- | --- | --- |
| *Coronaviridae* [ssRNA(+)] | 644,483 | 12 | 12 | Alpha, beta, gamma and unclassified coronaviruses |
| *Influenza* [ssRNA(-)] | 1,073,237 | 128 | 125 | Influenza A (H1N1, H2N2, H3N2, H5N1, H7N9, H9N2), Influenza B |
| *Metapneumovirus* [ssRNA(-)] | 443,974 | 5 | 1 | Metapneumovirus |
| *Rhinovirus* [ssRNA(+)] | 1,339,435 | 130 | 107 | Rhinovirus A and A1, B, C (C1, C2, C10), other enterovirus |
| SARS-CoV-2 [ssRNA(+)] | 865,303 | 87 | 1 | SARS-CoV-2 |
| *Human* | 3,531,425 | 1* | 1 | Human transcriptome |
| **Total** | **7,897,857** | **363** | **247** |  |

“\*”: GENCODE canonical transcripts were used to represent human reads in RNA-seq data.

### Model architecture

PACIFIC was implemented using the Keras API with a TensorFlow backend. Input reads were converted into 9-mers with a stride of 1, forming a vocabulary size of 4^9^ = 262,144 k-mers. Each of these k-mers is assigned a number using the Tokenize API from Keras (50) from 1 to 262,144. The first index position of 0 is reserved to denote zero-padding for variable length sequences. Tokens are fed into the first hidden layer of the neural network and transformed into continuous vectors of length 100. After the embedding, a convolutional layer takes the previous numerical vectors and uses 128 convolution filters with a kernel size of 3.

A pooling layer is used after the convolution, using *max pooling* with a kernel size of 3. A bidirectional long-short term memory (BiLSTM) layer then follows, which uses two traditional LSTMs; one starts ‘reading’ the input sequence from one of the two flanks, and the other from the opposite end. The output of the two LSTMs is then combined and passed to the next layer. Finally, PACIFIC has a fully connected layer using a *softmax* function to calculate posterior probabilities for each of the six classes. To reduce overfitting, we used 20% dropout at each hidden layer.

Cross-entropy was used as the loss function and ADAM (51) was used as the optimizer. The final configuration of the network, hyperparameter tuning and the number and configurations of layers was obtained after several iterations between training and validation data. The final model is implemented *as double prediction* on both strands of the input sequence, whereby the forward and reverse-complement of the input sequence are predicted for class assignment. Classes for both predictions were required to match. The threshold of posterior probability for the assigned class was ≥0.95.

### PACIFIC training

NVIDIA GeForce RTX2080Ti was used to accelerate training. We trained two LSTM implementations, one using the fast LSTM implementation backed by CuDNN, supported only with NVIDIA Graphical Processing Unit (GPU). The other model was built using the regular implementation of LSTM. Both models achieved the same results. We started the training by shuffling the training sequences, using chunks of 200,000 reads to avoid loading all reads into memory. 90% of the data was used for training and 10% for optimization of parameters. After 15 chunks, the model converged on the validation set and training was halted. During training, we used binary accuracy (1), categorical accuracy (2) and cross-entropy loss from the optimization set to monitor the training.

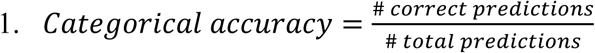

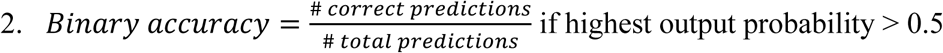

Training was completed when the model converged, obtaining final categorical and binary accuracy values of 0.99, and 0.003 for optimization loss.

### PACIFIC test datasets

We generated 100 independent test datasets using the ART sequence simulation software (version 2.5.8, (30)) with default error models for substitutions, insertions and deletions using the Illumina® HiSeq 2500 sequencing platform. For each dataset, we set seeds starting from 2021 to 2120 using a random number generator for reproducibility. Synthetic data contained 150nt single end reads derived from seven classes; the five model virus classes, a human class, and an “unrelated” class composed of 32,550 distinct virus genomes downloaded from the NCBI Assembly database. We sampled ∼100,000 reads per class using a class-specific fold-coverage parameter to generate ∼700,000 reads per test data (Table 5). Approximately 22% of reads contained mismatches, insertions or deletions relative to their respective reference sequences, reflecting error profiles of the Illumina sequencing platform. This process was automated using a custom script (generatebenchmarkdata.pl).

**Table 5.**
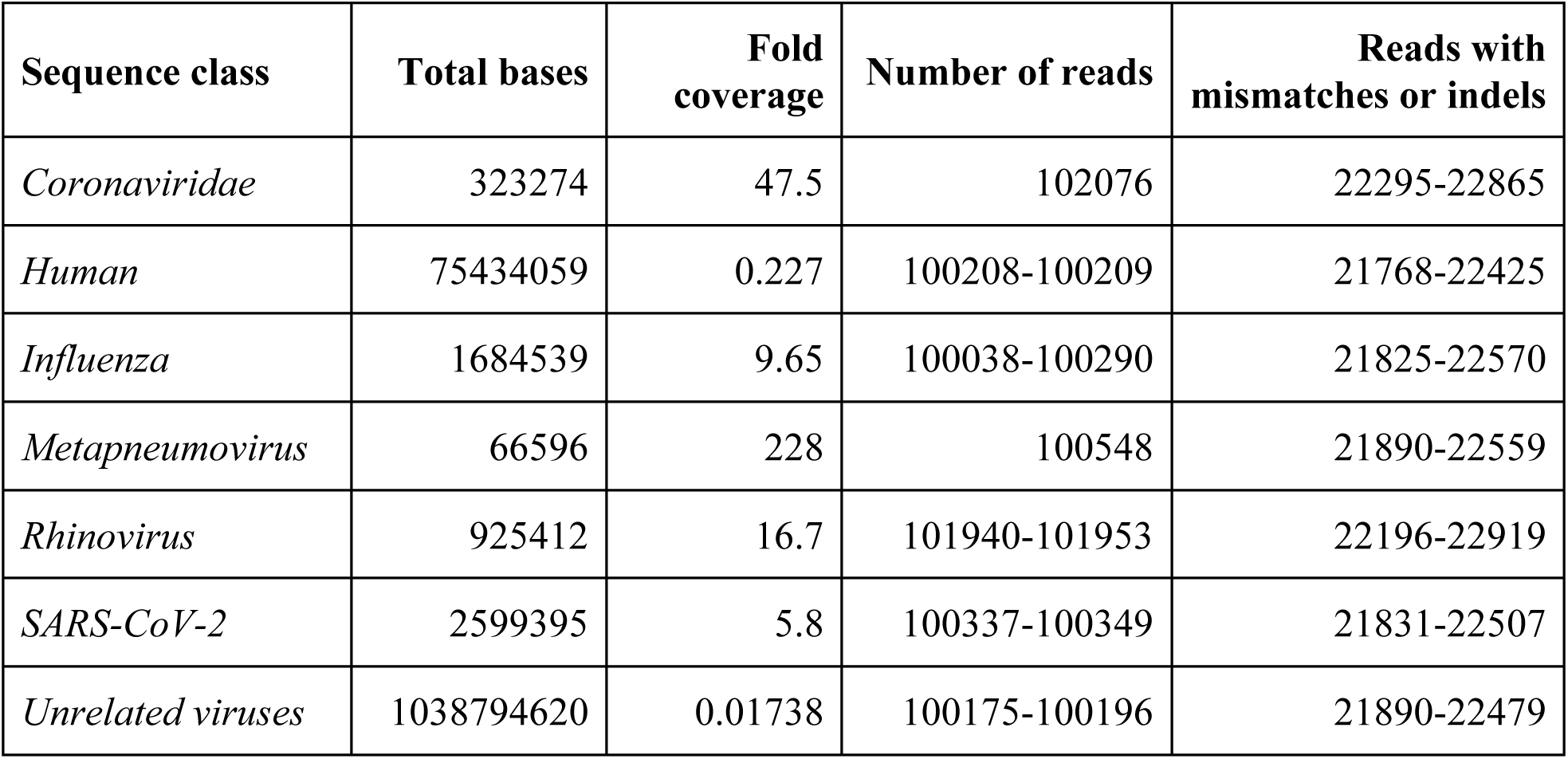
Summary of benchmark datasets

### PACIFIC performance tests

PACIFIC was used to assign class labels to reads in the test data, and performance metrics were calculated by comparing known and predicted labels for each read. A read was assigned a class if the maximum posterior probability score for a class was ≥0.95. A true positive (TP) was defined when the true label and the predicted label were the same for a read. A true negative (TN) was defined when a read that did not belong to the true class was correctly predicted as a class different from the true class. False positives (FP) were reads which were predicted to be as the true class, although they originated from a different class. False negatives (FN) were all reads belonging to the true class but were predicted as a different class. An example confusion matrix for SARS-CoV-2 is described in Table 6. Precision, recall, accuracy, false positive rate and false negative rate were calculated using equations 3-7 below.

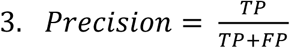

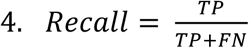

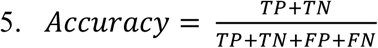

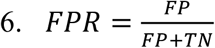

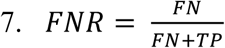

where TP = True positive, FP = False positive, TN = True negative, FN = False negative, FPR = False positive rate, FNR = False negative rate.

**Table 6.** Confusion matrix using SARS-CoV-2 as an example of a positive class.

|  | *True/ Actual condition* |  |  |
| --- | --- | --- | --- |
| *Predicted condition* |  | ***Positive***<br>*SARS-CoV-2 +* | ***Negative***<br>*All other classes* |
|  | *SARS-CoV-2 +* | True positive (TP) | False positive (FP) |
|  | *All other classes* | False negative (FN) | True negative (TN) |

### Establishing false positive rate thresholds for each class

This experiment was performed to quantify the impact of variable proportions of reads from each class on the percentage of false positives and to establish the detection threshold for each virus class in RNA-seq data. For each viral class in PACIFIC, we generated 100 datasets containing 500,000 reads derived from 4 out of the 5 viral classes, the human transcriptome and unrelated viral genomes in variable proportions. Reads were simulated using the ART software (30) with Illumina® HiSeq2500 error profiles that were 150nt long and modelled single end experiments. One of the five viral classes that was excluded was considered as the test class. This process was automated using a custom script (generatefprdata.pl).

Subsequently, PACIFIC was run in *double prediction* to assign classes to each read. To calculate the percentage of false positives in each experiment, we counted the number of reads predicted as the absent test class and divided by the total number of reads.

### Detecting viruses in human datasets and comparison with other tools

We downloaded 63 RNA-seq experiments from NCBI SRA database. Run accessions and other metadata details are supplied in Additional file 1: Table S2. All data were downloaded from the NCBI database using the SRA Toolkit *prefetch* and *fastq-dump* commands and applying the *--gzip* and *--fasta* options (52). For the GALA II cohort study with 48 RNA-seq datasets and read lengths 18-390nt, we discarded reads <150nt long. We then used PACIFIC to assign the presence/absence of each virus class in all 63 samples using the detection thresholds established in the previous section. We compared PACIFIC’s predictions with two alternative methods for virus detection: an alignment-based approach using BWA-MEM (53), and a k-mer based approach using Kraken2 (19), described below.

For BWA-MEM (53), all reads were mapped using default parameters to a combined reference containing assembly sequences for the five viral classes and the human transcriptome used for training PACIFIC (Table 4). Reads were assigned to a virus class based on the class membership of the genome assembly as described in Table 4 and Additional file 1: Table S1. For Kraken2, we first downloaded the Kraken taxonomy database and built a k-mer database using the same genomes used to train PACIFIC (Table 4). Kraken2 was then run using the – *use-names* flag, and output reads were parsed using species scientific names and reads were assigned a class based on the class membership of the genome assembly (Additional file 1: Table S1, Table 4). To fairly compare all three methods, we applied class detection thresholds as determined for and used in PACIFIC (Table 2) for the presence or absence of a virus class within a sample.

To investigate the origin of reads for all reads in samples that were discordantly predicted for the presence of a virus class by PACIFIC, BWA-MEM or Kraken2, we used the BLAST suite (v2.10.1+) (54,55) to align reads to the NCBI nucleotide (*nt*) database, which includes sequences from all domains of life. We took the best hit from the pairwise alignment for each read, filtering for alignments with an E-value <1e-6. BLASTN was used with the following parameters: *-task ‘megablast’ -max_target_seqs 1 -max_hsps 1 -evalue 1e-6* to query discordant viral class assignments between PACIFIC, BWA-MEM and Kraken2.

## Declarations

### Ethics approval and consent to participate

Not applicable.

### Consent for publication

Not applicable.

### Availability of data and materials

Source code is available in the PACIFIC Github repository [https://github.com/pacific-2020/pacific/] under the MIT License. Publicly available data generated or analysed during this study are included in the following published articles (31,32,34), accession numbers and other metadata are described in [https://github.com/pacific-2020/pacific/tree/master/metadata]. Other supplementary information and test data are available and can be downloaded from [https://cloudstor.aarnet.edu.au/plus/s/sRLwF3IJQ12pNGQ.

### Authors’ contributions

PAM, RFB, EE and HRP conceptualised the project. PAM contributed to the training and initial testing of the model. PAM, RFB and HRP performed all data analyses with input from EE. HRP, SE and EE supervised the project. PAM, RFB, EE and HRP drafted the manuscript. All authors read and approved the final manuscript.

## Acknowledgements

The authors would like to dedicate this study to Smt. Mukta Sheladiya and all other unfortunate souls who lost their battle against the COVID-19 infection. We would like to offer our sincere gratitude to all healthcare workers who have supported millions of people during these trying times across the world. We thank Dr. Cheng Soon Ong for his input in the development of the model. Finally, we thank Dr. Saul Newman and Dr. Teresa (Terry) Neeman for their valuable feedback and comments in preparation of the manuscript.

## Supplementary Information

**Additional file 1.** Supplementary Tables S1, S2, and S3.

- **Supplementary Table S1**: Summary table of genomes and assemblies used to train PACIFIC.
- **Supplementary Table S2**: PACIFIC testing metrics.
- **Supplementary Table S3.** Publicly available samples used to run PACIFIC, BWA and Kraken2.

**Additional file 2.** Supplementary Figures S1 and S2, Details of the BLAST analysis.

