## Supplementary Figures for "PACIFIC: A lightweight deep-learning classifier of SARS-CoV-2 and co-infecting RNA viruses"

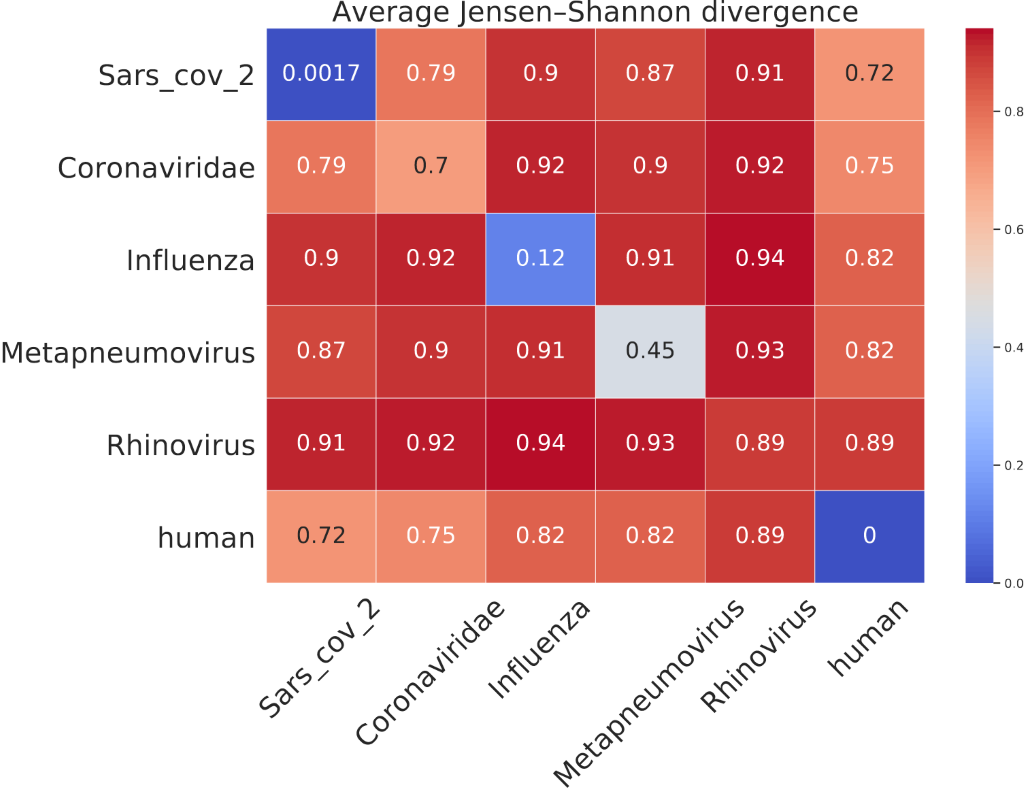
**Supplementary Figure 1.** Inter/intra-class Jensen-Shannon Divergence (JSD). All-vs-all JSD were calculated using 9-mers and averages calculated at a class level. Pair of identical pairs have a JSD of 0 for the pair (blue), while sequences that do not share any k-mers have a JSD value of 1 (red). Intra-class JSD were overall lower compared to the inter-class JSD as expected.

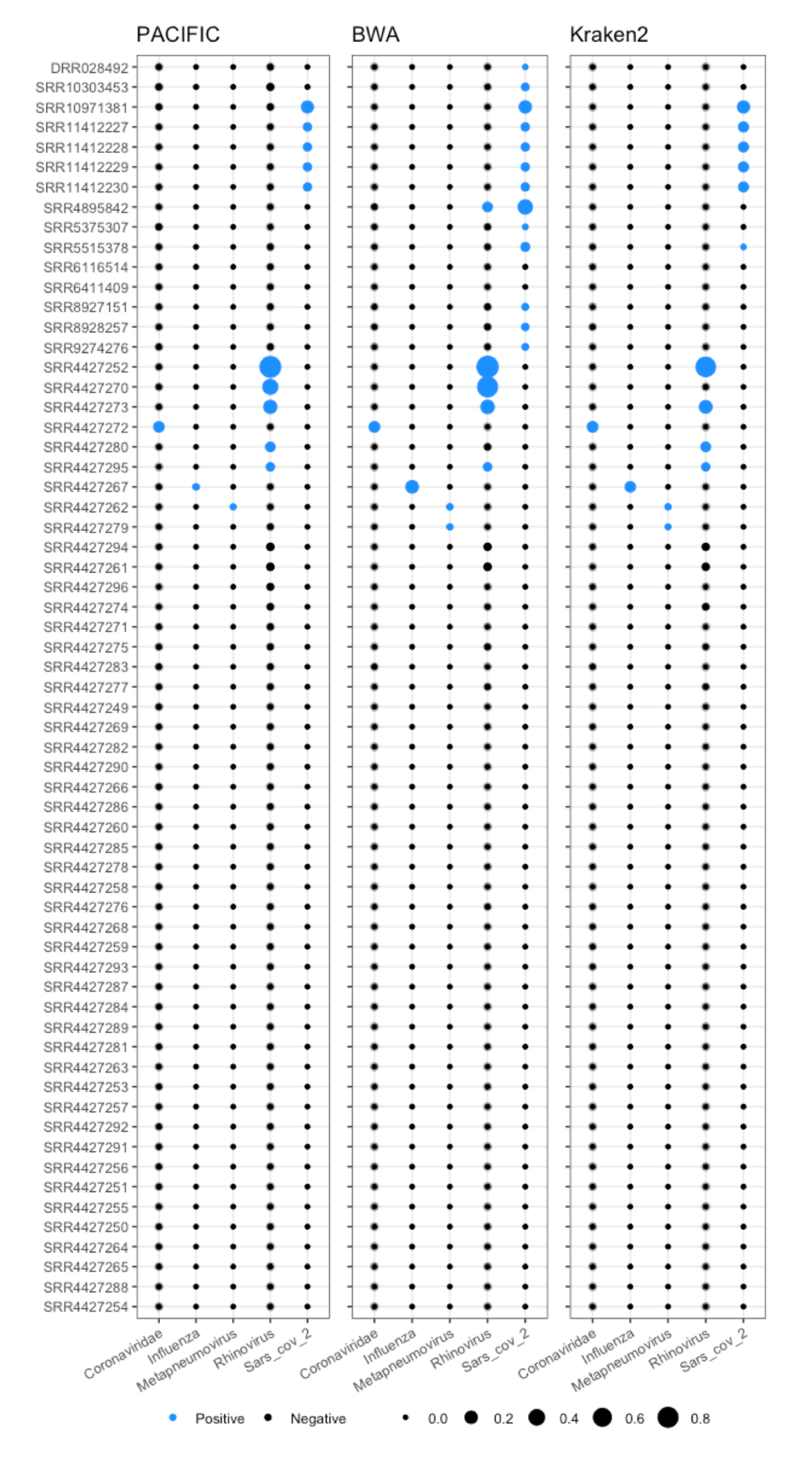

**Supplementary Figure 2.** PACIFIC, BWA-MEM and Kraken2 predictions on all samples from all datasets used in this study (31,32,34). Spheres represent the percentage of predicted reads above 0.95 posterior probability score for PACIFIC (left), and the proportion of predicted viral reads using BWA-MEM (centre) and Kraken2 (right). Black circles represent predictions where the percentage of reads does not reach PACIFIC thresholds (Table 2). Blue circles represent samples predicted above PACIFIC class thresholds.

*Extension of BLAST analysis*

One sample (SRR4427279) was classified as being positive for the Metapneumovirus class by BWA-MEM and Kraken2 but not by PACIFIC. Best hits for all 28 reads aligned to human respiratory syncytial virus A (see Results). To further test whether these reads may be derived from metapneumovirus, we investigated secondary BLAST hits for these reads. We used BLASTN to align all 28 reads to the NCBI nucleotide *(nt)* database, taking the best hsp from pairwise alignments for a maximum of 10 target sequences for each read, using the parameters --*max_target_seqs 10 –*max_hsps *1*. Such searches revealed that all 28 reads had their best hits for their top 10 alignments to either human respiratory syncytial virus A sequences (267/280 pairwise alignments) or its genus Human orthopneumovirus (13/280 pairwise alignments); no hits to metapneumovirus sequences were found.

BWA-MEM, but not PACIFIC or Kraken2 also reported reads that aligned to Rhinovirus for one sample, with best hits from BLAST showing alignments primarily to *Homo sapiens* and *Pan paniscus* (SRR4895842; see Results). A similar strategy was used to test whether secondary hits from these reads were derived from rhinovirus using BLAST. We collected reads classified as Rhinovirus by BWA-MEM and used BLASTN against (*nt*) database to investigate the 10 best database matches per read. The majority of these hits belong to *Homo sapiens*, followed by *Pan paniscus.* In total, counting best and secondary hits there were 52 taxonomy IDs matching to these reads; none of these IDs corresponded to Rhinovirus, or any of the other model virus classes.
